# Behavioral and imaging analysis of Foxg1 heterozygous mice

**DOI:** 10.1101/2022.03.29.486318

**Authors:** Kirsty R. Erickson, Rebekah Lifer, Jonathan K. Merritt, Zeljka Miletic Lanaghan, Mark D. Does, Karthik Ramadass, Bennett A. Landman, Laurie E. Cutting, Jeffrey L. Neul

**Author notes:** **Author Contributions** Conceptualization, Methodology, and Funding Acquisition: JLN; Investigation: KRE, RL, JM, ZML; Formal Analysis: KE, JKM, JLN, MDD, KR, BAL, LEC; Writing-Original Draft Preparation: JLN; Writing-Review & Editing: JLN, KRE, RL, JKM, ZML, MDD, KR, BAL, LEC, JLN.

## Abstract

FOXG1 Syndrome (FS) is a devastating neurodevelopmental disorder that is caused by a heterozygous loss-of-function (LOF) mutation of the *FOXG1* gene, which encodes a transcriptional regulator important for telencephalic brain development. People with FS have marked developmental delays, impaired ambulation, movement disorders, seizures, and behavior abnormalities including autistic features. Current therapeutic approaches are entirely symptomatic, however the ability to rescue phenotypes in mouse models of other genetic neurodevelopmental disorders such as Rett syndrome, Angelman syndrome, and Phelan-McDermid syndrome by postnatal expression of gene products has led to hope that similar approaches could help modify the disease course in other neurodevelopmental disorders such as FS. While FoxG1 protein function plays a critical role in embryonic brain development, the ongoing adult expression of FoxG1 and behavioral phenotypes that present when FoxG1 function is removed postnatally provides support for opportunity for improvement with postnatal treatment. Here we generated a new mouse allele of *Foxg1* that disrupts protein expression and characterized the behavioral and structural brain phenotypes in heterozygous mutant animals. These mutant animals display changes in locomotor behavior, gait, anxiety, social interaction, aggression, and learning and memory compared to littermate controls. Additionally, they have structural brain abnormalities reminiscent of people with FS. This information provides the framework for future studies to evaluate the potential for post-natal expression of FoxG1 to modify the disease course in this severe neurodevelopmental disorder.

## Introduction

FOXG1 Syndrome (FS, OMIM # 164874), previously considered a congenital variant of Rett syndrome, is a devastating neurodevelopmental disorder caused by a heterozygous loss-of-function mutations of the *FOXG1* gene (1, 2). Nearly all cases of FS are caused by *de novo* mutations in *FOXG1* (3), which encodes a transcriptional repressor, forkhead box G1 (FoxG1) that plays an important role in telencephalic development (4). People with FS have severe developmental delay and fail to gain many skills, movement disorders including dyskinesia, seizures, difficulty, or lack of the ability to independently ambulate, seizures, and autistic features (3). Neuroanatomical features include corpus callosum agenesis, pachygyria, postnatal microcephaly, and moderate-to-severe myelination delay (5).

Currently, therapies for FS are entirely symptomatically based and do not alter the overall course of disease to any significant degree. Affected individuals have markedly decreased quality of life and require full-time care for activities of daily living. One challenge in the development of disease modifying therapies in FS is the fact that the diagnosis is made after birth, after the altered embryonic brain development resulting in the observed structural brain abnormalities. Although FoxG1 function plays a key role in embryonic brain development, FoxG1 continues to be expressed in the postnatal brain. Support for an ongoing postnatal role of FoxG1 has been demonstrated by experiments removing FoxG1 function postnatally from specific brain regions (6) or neuronal cell types (7), resulting in disruption of the structure of the dentate gyrus of the hippocampus (6) or alterations in learning and memory and social behavior (7). These findings provide hope that restoration of FoxG1 function, even postnatally, could be disease modifying. Recent work on mouse models of other neurodevelopmental disorders, such as Rett syndrome (RTT), have demonstrated the possibility of a reversal of disease via adult post-symptomatic re-expression of *MECP2*, the gene disrupted in RTT (8). Similarly, postnatal restoration of correct gene expression levels has been shown to be beneficial in mouse models of other neurodevelopmental disorders, such as MECP2 Duplication Syndrome (9), Angelman syndrome (10–12), and Phelan-McDermid Syndrome (13), raising the question whether a similar opportunity exists for modification of symptoms of FS with postnatal expression of FOXG1. Encouragingly, recent work demonstrated that early life transplantation of GABAergic neuronal precursors could improve some phenotypes (14).

A current issue related to the development of such therapies in FS is the limited evaluation of phenotypes in mice with heterozygous mutations in *Foxg1*, as most work has focused on the early developmental effects of homozygous loss. Reduction in volume in the neocortex, hippocampus, and striatum have been seen in heterozygous *Foxg1* mutant mice (15). Behavioral analysis found changes in locomotor activity and memory in one mutant line (16), and altered social behavior, poor working memory, and decreased anxiety in a different mutant line (14). Additionally, neurophysiological experiments have found alterations in EEG power spectral features (14) and visual evoked potentials (17). Here we present the generation and initial characterization of a new mouse model of FS, which was generated by the insertion of a loxP-flanked “STOP” cassette containing a splice acceptor to disrupt translation of the FoxG1 protein. This resulted in a decrease in FoxG1 protein expression and alterations in locomotion, gait, learning and memory, social behavior, and overall and regional brain volumes. This work provides the framework for future experiments to evaluate the overall ability of restoration of FoxG1 at various developmental timepoints to modify important clinically relevant phenotypes.

## Materials and Methods

### Mouse care and model generation information

All methods and animal care procedures were approved by the Vanderbilt Animal Care and Use Committee (Protocol Number: M1700069-01), and all aspects of the study were carried out in accordance with the recommendations in the Guide for the Care and Use of Laboratory Animals of the National Institutes of Health. Mice were housed in AAALAC-approved facilities at Vanderbilt University Medical Center. Euthanasia was performed via overdose of inhaled anesthetic agent (isoflurane) followed by decapitation and removal of vital organs. Heterozygous *Foxg1*^*tm4144Tac*^ (referred to as “MUT”, description of engineering to create this allele of *Foxg1* provided below) were mated to wild type C57BL/6J mice to generate experimental animals: *FoxG1*^*MUT/WT*^ heterozygous mice and wild-type littermates which were used as controls (WT). Both male and female animals were used.

*Foxg1*^*tm4144Tac*^ was generated by Taconic Biosciences using a targeting strategy designed to insert a loxP-flanked transcriptional termination cassette (STOP) into intron 1 as well as adding 3xHA-tag on the carboxy-terminus of *Foxg1*. The STOP cassette contains a splice acceptor (SA), a combination of polyadenylation signals (human Growth Hormone and synthetic polyadenylation signals) and translation termination codons in all three reading frames. The sequence for the 3xHA-tag was inserted in-frame between the last amino acid codon and the translation termination codon in exon 2. The targeting vector included a positive selection marker (Puromycin resistance) flanked by FRT sites and inserted into the loxP-flanked STOP cassette. Homologous recombinant clones were generated in the Taconic Biosciences C57BL/6N Tac ES cell lines with positive (PuroR) and negative (Thymidine kinase) selection. Chimeric animals (G0) were generated and degree of chimerism (as judged by coat color contribution of the ES versus BALB/c host), and highly chimeric animals bred to C57BL/6N-Tg(CAG-Flpe)2Arte animals to remove the FRT-flanked selection cassette (See Fig. 1A for diagram of the engineered allele after removal of the FRT selection cassette). One chimera with high degree of chimerism generated 3 germline transmitted offspring which were used to establish *Foxg1*^*tm4144Tac*^ line which was transferred to the Neul lab and subsequently backcrossed to C57BL/6J for >5 generations. PCR genotyping was performed using the following primer sets: Mutant allele (326bp band size):

**Figure 1:**
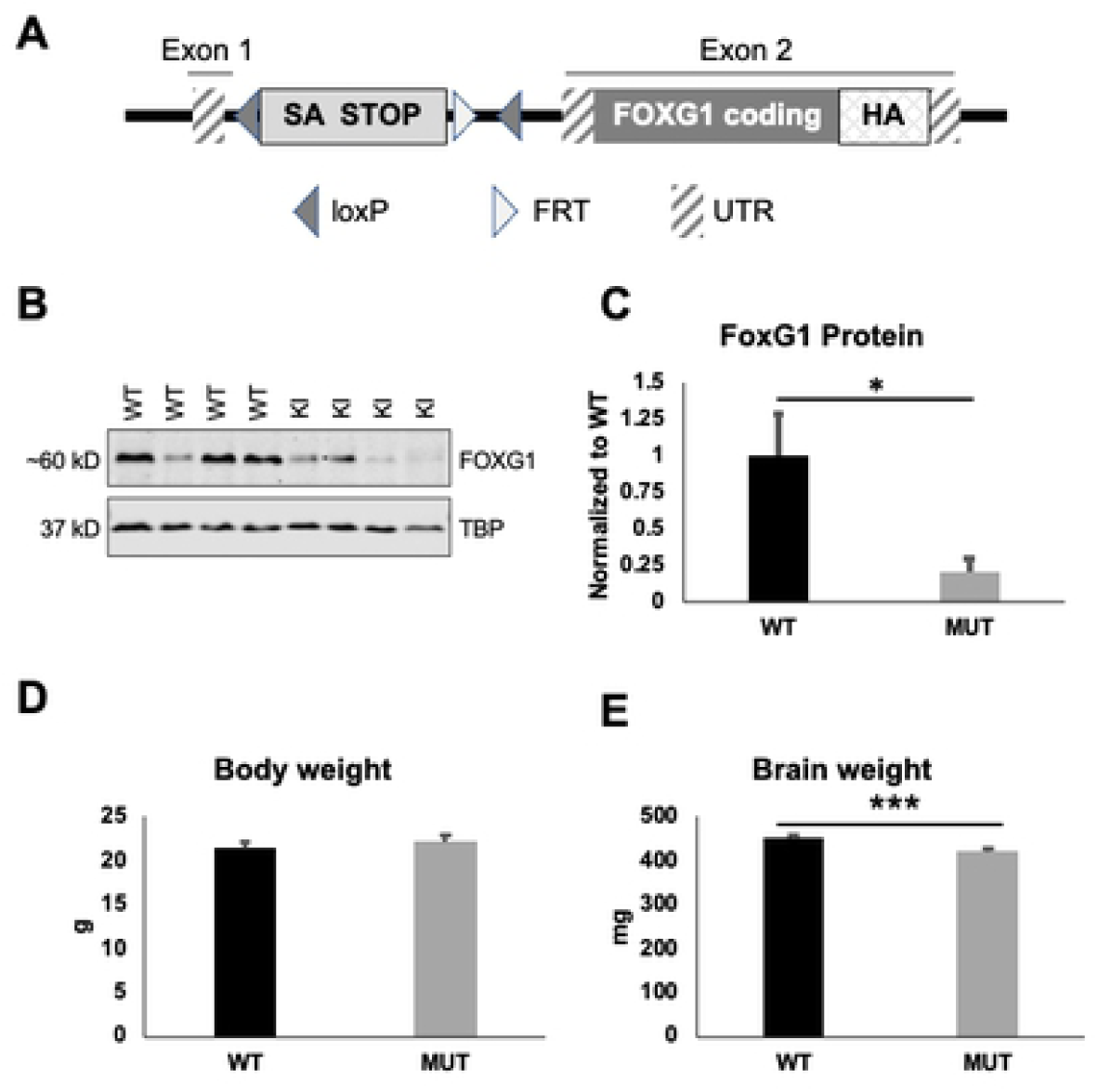
Generation of new allele of *Foxg1*. A) Design of the engineered allele of *Foxg1*. Endogenous exons are indicated above. loxP sites are identified by left oriented triangle and FRT site by right facing triangle. Untranslated regions (UTR) are designated with diagonal stripes, endogenous coding sequence with dark grey, and inserted STOP cassette containing a splice acceptor (SA) designated in light grey. Three HA tags were inserted in-frame at the carboxy-terminus. B) Western blot of FoxG1 protein in brains from heterozygous *FoxG1*^*MUT/WT*^ (MUT) animals compared to *FoxG1*^*WT/WT*^ (WT) animals. C) Quantification of FoxG1 protein from WT and MUT animals, normalized to average WT protein amount. D) No difference in body weight between WT and MUT animals [F(1,28)=0.417, p=0.523]. E) MUT animals have decreased brain weight compared to WT animals [F(1,28)=19.174, p<0.0001). For panels D, E age =16-17 weeks, WT n=15, MUT n=15. No significant genotype*sex interaction observed. *p<0.05, **p<0.01, ***p<0.001

Forward primer (13959_32): GATAAATATTGACGCGCAAAGG

Reverse primer (13959_33): TGTGCTGGTACTGTTTCTGAGC

Wild type allele (585bp)

Forward primer (1260_1): GAGACTCTGGCTACTCATCC

Reverse primer (1260_2): CCTTCAGCAAGAGCTGGGGAC

### Physical post-mortem characterization

At 16-17 weeks of life mice were humanly euthanized and whole body and dissected brain weights obtained.

### Western Blotting

Protein lysates for western blotting were prepared by homogenizing whole brains isolated from e16.5 *FoxG1*^*MUT/WT*^ embryos in Tris lysis buffer (50 mM Tris-HCl pH 7.5, 150 mM NaCl, 1% NP-40, 1 mM EDTA, 1 mM EGTA, 1 mM DTT, 1 μM Pepstatin, 10 μM Leupeptin, 200 μM PMSF). Protein concentration was determined using the 660nm protein assay (Pierce 22660). 5 ug of total protein for each sample was separated on a 10% SDS-PAGE gel and transferred to a nitrocellulose membrane using standard methods. Following western transfer, membranes were incubated with primary antibodies (1:1000 rabbit anti-Foxg1 (Abcam ab18259) and 1:5000 mouse anti-TBP (Abcam ab51841)) overnight at 4 degrees C. The following day, membranes were incubated with secondary antibodies (1:10,000 goat anti-rabbit 800CW (Licor 926-32211) and 1:10,000 goat anti-mouse 680RD (Licor 926-68070)) for 1 hour prior to imaging on a Licor Odyssey CLX. Quantification was performed using Licor Image Studio.

### Behavioral characterization

All behavioral experiments were performed in the Vanderbilt University Neurobehavioral Core Facility. Behavioral tests were performed using male and female mice. Mice were group housed (3-5 mice/cage) on a 12-hour light/dark cycle with food and water available ad libitum. Mice were transferred to test rooms and acclimated to the environment for 30 minutes prior to testing. Unless otherwise stated, all equipment was cleaned with 70% ethanol between trials to provide a standardized testing environment. All behavioral experiments were performed during the light phase. All data is provided in Supplemental Excel Sheet.

#### Rotarod

Motor coordination and learning was assessed using a five-lane MedAssociates accelerating rotarod, similar to methods described previously (18). Briefly, mice were placed on a textured cylinder (3.18cm in diameter) located 30cm above the apparatus floor. Once all the mice were loaded, the rotarod was set to initiate preprogrammed acceleration from 4-40rpm at 4rpm. The latency to fall was recorded, with a maximum time of 300 seconds. To account for passivity in the task, if an animal completed two passive rotations (i.e. underwent two full rotations without making a step forward) it was considered to have fallen and the time was recorded as the latency to fall time. Animals underwent three trials a day for three consecutive days, with at least a 30-minute intertrial interval. Time to fall was averaged across the three trials for each day.

#### Open field analysis

Exploratory locomotor activity was assessed in chambers measuring 27 × 27 × 20.5 (MedAssociates), housed in sound-attenuating cases over a 30-minute period using the method described previously (19). Activity was captured via infrared beams and detectors. A region around the exterior perimeter of the chamber was designated as the “surround”, and the inner portion is detected as the “center”. General activity was indicated by the time and distance traveled. The proportion of time and distance in the center relative to time or distance in surround regions of the chamber were evaluated.

#### Elevated Zero Maze

Anxiety-like phenotypes were assessed in an elevated zero maze (Stoelting: 50cm inner diameter, 5cm lane width, 15cm closed arm wall heigh, and 50cm apparatus height), as previously described (19). Illuminance was measured as approximately 300 lux in the open arm and 70 lux in the closed arm. In this task, the mice were placed in the open arm of the maze at the beginning of each trial and behavior was assessed for 5 minutes. Video data was analyzed by ANY-maze software (Stoelting). Following the test mice were placed into a clean cage, to avoid any confounding effect on naïve mice.

#### Forced Gait Analysis

Gait was assessed using the CleverSys Treadscan for forced gait analysis, using a method based on that described previously (19). The apparatus consists of a transparent treadmill with a high-speed digital camera under the treadmill that captures foot placement of the mice on the treadmill. Mice were allowed to habituate to the treadmill chamber for 1 minute prior to the start of the task. The treadmill was then turned on and off (20 cm/s) until the test mouse was continuously running. A 20 second video was then captured. CleverSys analysis software was used to identify periods of continual running for analysis. These video segments were then subject to footprint analysis with paw identification training modules in accordance with the manufacturer’s instructions. The data fields from the manufacturer’s output were simplified by averaging left and right results together from either forepaws or hindpaws to get a single value. For example, Right Forepaw Stance time and Left Forepaw Stance time were averaged to calculate Forepaw Stance time for each animal. Because many of the parameters contained in the output are related to each other, we further streamlined the data fields to be analyzed by generating a correlation matrix with all data present (ignoring sex and genotype). A set of 22 variables were selected that were not strongly correlated and represented the highly correlated data fields excluded.

Descriptions of parameters assessed is included below:

The stance time is the amount of time elapsed while the foot is in contact with the runway. The brake time is the time elapsed between the start of a stance and the instance the foot reaches the normal stance position of the front feet, when forced is applied to move the body. The homologous coupling is the fraction of the stride of a reference foot, where the given foot on the same half starts its stride. It is the same as the coordination between left and right foot on the same girdle. The body rotation is the average orientation direction measured in degrees and measures the overall orientation of the animal i.e., the orientation of the body while the animal is walking.

#### Fear Conditioning

The fear conditioning assay was used to test learning and memory, similar to the method outlined previously (20). On training day, mice were acclimated in a room adjacent to the test room for 30 minutes. After acclimation, each mouse was moved to the test room and placed into a 29.53 × 23.5 × 20.96 cm MedAssociates chamber equipped with a stainless-steel grid floor for delivery of an electric shock. Mice moved within the chamber for 2 minutes, and then a 30 second, 80dB white noise tone was then administered with the last 2 seconds accompanied by the delivery of a 0.5mA foot shock. This tone-shock pairing was repeated after a 2-minute interval, after which mice were left in the chamber for 1 minute and then removed. Mice were then moved to a fresh cage in an adjacent recovery room to avoid confounding effects with naïve cage-mates. Twenty-four hours later, mice were tested for context-dependent fear memory by placing them back into the testing chamber for 4 minutes. After two hours mice were assessed for cue-dependent fear memory. Mice were placed into an altered testing environment for 4 minutes, during which the final 2 minutes were accompanied by the 80dB tone stimulus. The altered environment consisted of red lighting in the acclimation/test/recovery rooms, lack of white light in the chamber, a flat baseboard, rounded walls, and vanilla scent. Video cameras mounted on the front wall of the testing apparatus recorded the movement of each mouse, and freezing behavior was assessed during the training period, across the full 4-minute context test, and the final two minutes of the cue-test. Freezing was defined as behavior below a motion threshold below 18 arbitrary units for 30 frames (1 second) minimum freeze during, and the percent of time freezing was calculated using the default linear method.

#### Marble Burying

Mice were placed in individual home-like cages containing 1-inch of bedding, that had been smoothed and slightly compacted, along with 20 dark blue marbles arranged into 5 rows of 4 marbles. The mice were allowed 30 minutes to investigate the marbles. At the end of the task the mice were removed and placed back into their home-cage, and the number of marbles buried at least 50% into the bedding was scored and recorded manually.

#### Tube Test for Social Dominance

To test for social interaction phenotypes we assessed mice using the tube test for social dominance. Test mice and a sex-, weight-, and age-matched unfamiliar conspecific are placed on opposite ends of a ∼3 cm square and 20 cm long plexiglass tube. The duration of the test and which mouse backs out first was recorded.

#### Nest Building

Nest building is an innate behavior in rodents and was used to assess the general well-being of *Foxg1* Het mice as well as for impaired activities in daily living. The procedure used was based on previous work (21). On the afternoon of test day, mice were placed into clean individual cages containing pre-weighted cotton nestlets (5 × 5 × 0.3 cm, ∼2g) in the middle of the cage. The cages were placed into an environmentally controlled chamber overnight, and the nest quality and weight of the remaining nestlet were made the next morning (approximately 12 hours). Nest quality was scored using the method outlined previously (22).

#### 3-Chamber Social Interaction Test

Social behavior was assessed using the Crawley 3-chamber assay, using the method described previously (19). The apparatus consists of a clear 60 × 42 × 22 cm box, divided into three adjacent and equally sized compartments. Openings within the walls that separate the compartments allow for the mice to travel freely between them. Empty inverted wire pencil cups were placed in the same-sided corners of each left and right compartment. There were three separate stages of testing conducted in a single day. The first stage, habituation, allowed the mice to freely explore the apparatus for 5 minutes. In stage two, sociability, an unfamiliar sex-, age-, and weight-matched conspecific (Stranger 1) was placed in one of the pencil cups. Test mice were reintroduced to the apparatus for 10 minutes. Stage three, social novelty preference, consisted of another unfamiliar sex-, age, and weight-matched mouse (Stranger 2) was placed under the remaining cup. Test mice were again allowed to explore the apparatus for 10 minutes. Stranger 1 and Stranger 2 were always from different cages and had no previous contact with experimental mice. The stranger mice were also habituated for thirty minutes a day for two days prior to testing. After stage three, test mice were placed into a clean cage to prevent contact with untested cage mates. A camera mounted above the testing apparatus captured videos for all stages, and then manually scored for interaction with the pencil cups during stages two and three. We defined interaction as sniffing, pawing, or rearing onto the cup. Discrimination indices were calculated for sociability and social novelty, stage 2 (Sociability) and stage 3 (Social Novelty) as described below:

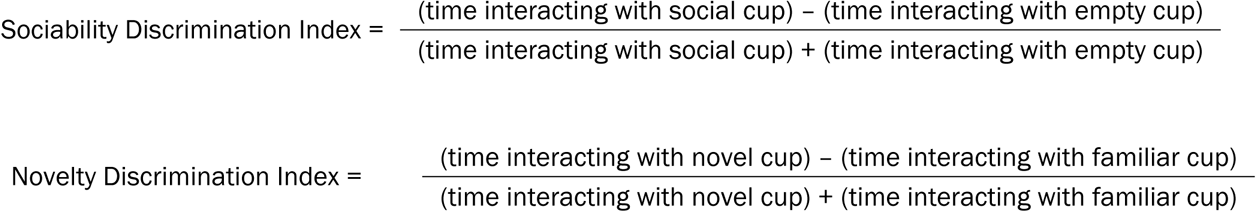

### Brain Imaging

#### Extraction of Brains

Each mouse was deeply anesthetized with isoflurane and perfused with 60mL 1X PBS at 6mL/min to flush out body fluid. Next, they were perfused with 40mL of fixative (2% PFA, 2.5% glutaraldehyde, 1mM gadolinium, 1X PBS) at 6mL/min. Brain extraction was performed immediately after perfusion and immersed in 40mL of fixative for 1 week. After 1 week, the brains were moved to 25mL of a solution containing 1X PBS, 1mM gadolinium, 0.01% sodium azide, with this solution changed 4 times during a 1-week period. Animals were approximately 48 weeks old at the time of brain extraction, 6 animals for each genotype were used.

#### Imaging Acquisition

Mouse brain imaging was performed on a 7T 16-cm bore magnet operated by a Bruker Biospec console (Billerica, MA, USA) using a 25mm ID Doty RF litzcage coil (Doty Scientific, Columbia, SC). Four mouse brains were scanned per session and three MRI scan types were acquired: high resolution anatomical (HRANAT), selective inversion recovery, and diffusion tensor imaging. Total overall scan time per session was ≈ 15 h. Volumetric findings were the focus of the current study; therefore, only HRANAT parameters are detailed. HRANAT imaging data were acquired using a 3D RARE (rapid acquisition with refocused echoes) scan with repetition time (TR) = 350 ms, echo time (TE) and echo spacing (ESP) = 14 ms, echo train length (ETL) = 4, and 2 signal averages. Non-selective excitation and refocusing hard pulses of 200 µs and 125 µs were used respectively. Receiver bandwidth (BW) for signal acquisition = 50 kHz. Images were acquired with a field-of-view (FOV) = 21.6 × 16.2 × 14.4 mm3 and matrix size = 432 × 324 × 288 for nominal isotropic resolution of 50 × 50 × 50 µm3.

#### Imaging Analysis

All volumes were converted to nifti data format. For each structural volume, the 2006 Mouse Minimum Deformation Atlas (MDA, https://resource.loni.usc.edu/resources/atlases-downloads/) was registered to the acquired volume with a 12 degree of freedom affine transform using FSL Flirt version 6.0, and then the associated label file was transformed with nearest neighbor interpolation. Next the atlas intensity image was nonlinearly warped to the affine coregistered target using ANTs SyN (3.0.0.0.dev62-g1904a, http://stnava.github.io/ANTs/) with the following parameters: interpolation=Linear, winsorize-image-intensities=[0.005,0.995], use-histogram-matching=0, transform=SyN[0.1,3,0], metric =CC, convergence=[100×70×50×20,1e-6,10], shrink-factors=8×4×2×1, and smoothing-sigmas 3×2×1×0vox. The label files were transformed with the same transform using ANTs multi-label interpolation. Volumes were computed based on the total number of voxels of each label class in the final transformed image multiplied by the dimensions of the image. In all, the Atlas produced 23 Regions of Interest (ROIs, see Table X). Within these 23 ROIs, nine were hypothesized to be significantly smaller in volume for HET versus WT mice, eight were hypothesized to not be significantly different between mouse type. For the remaining volumes, we did not have specific hypotheses.

In addition to the automated image analysis outlined above, manual tracing of the corpus callosum/external capsule was performed by hand tracing this ROI from each slice. The total number of voxels for the ROI was multiplied by the image resolution (150×150×150 micrometers) to get the total volume.

### Statistical Analysis

For the total weight, brain weight, and all behavioral tests, two-way ANOVA (factors genotype, sex, with interaction term genotype*sex) was performed using SPSS version 25 (IBM Corp, 2017). We did not observe any significant genotype*sex interaction effects so subsequent analysis was performed using one-way ANOVA (factor genotype) and reported throughout. All data was included in analyses. Plots were generated in Excel (Microsoft) and display mean ± standard error of the mean. Statistical significance is represented in all plots as follows: *p<0.05, **p<0.01, ***p<0.001, ****p<0.0001.

For imaging analysis, Whole brain volumes (WBVs) and each of the 23 ROIs from the automated analysis was evaluated for group differences using t-tests (unequal variances assumed) in SPSS version 25 (IBM Corp, 2017). To avoid Type I error, False Discovery Rate (FDR) corrections were implemented to correct for multiple comparisons. FDR-corrections revealed that a *p* < .007 was required to reach statistical significance for the analyses with the WBV and 23 ROIs, while a *p* < .008 was required for the analyses on the 23 WBV-corrected ROIs. The manual tracing ROI was analyzed using independent samples t-test.

## Results

### Generation of new mouse allele of Foxg1

We generated a new allele of *Foxg1* using a commercial vendor by inserting a loxP-flanked STOP cassette containing a splice acceptor (SA) and a combination of polyadenylation signals and translation termination codons in all three reading frames into intron 1 (Fig. 1A). This is predicted to prevent translation of the FoxG1 protein which is entirely contained in exon 2. Western blotting of FoxG1 protein from E16.5 brains from WT or MUT animals (Fig. 1B, see Supplemental Data S1_raw_images for uncropped Western Blot images) found that MUT animals expressed ∼28% of WT FoxG1 protein levels (Fig. 1C). We did not observe any difference in overall body weight in MUT animals (Fig. 1D), however MUT animals had decreased brain weight compared to WT littermate controls (Fig. 1E).

### Heterozygous FoxG1 mice have alterations in locomotor activity

Because people with FS have a variety of movement abnormalities, we characterized locomotor coordination of male and female *FoxG1*^*MUT/WT*^ (MUT) using the accelerating rotating rod and found no difference in locomotor coordination on Day 1 or any changes in locomotor learning over the three-day test period, as measured by the average fall time on each day, compared to wild-type littermate controls (WT, Figure 2A). When overall locomotor activity was assessed using the Open Field Assay (OFA), MUT mice traveled less overall distance (Figure 2B) and had less vertical movements (Figure 2C). Additionally, MUT mice spent less percentage time and traveled less percentage distance in the center of the open field (Figure 2C, D). Because decreased time and distance in the center chamber of the open field can be interpreted as increased anxiety, we tested anxiety on these mice using the Elevated Zero Maze (EZM). There was no difference between MUT and WT animals in the distance travelled in the EZM (F[1,23]=0.299, p=0.590). MUT animals did not show increased time in the open arms of the EZM (Figure 2F) but did have an increased percentage of distance travelled in the open arms (Figure 2G), indicating decreased anxiety in contrast to the suggestion of increased anxiety observed in the OFA. Details of ages and number of animals for all tests presented in the figure legend.

**Figure 2:**
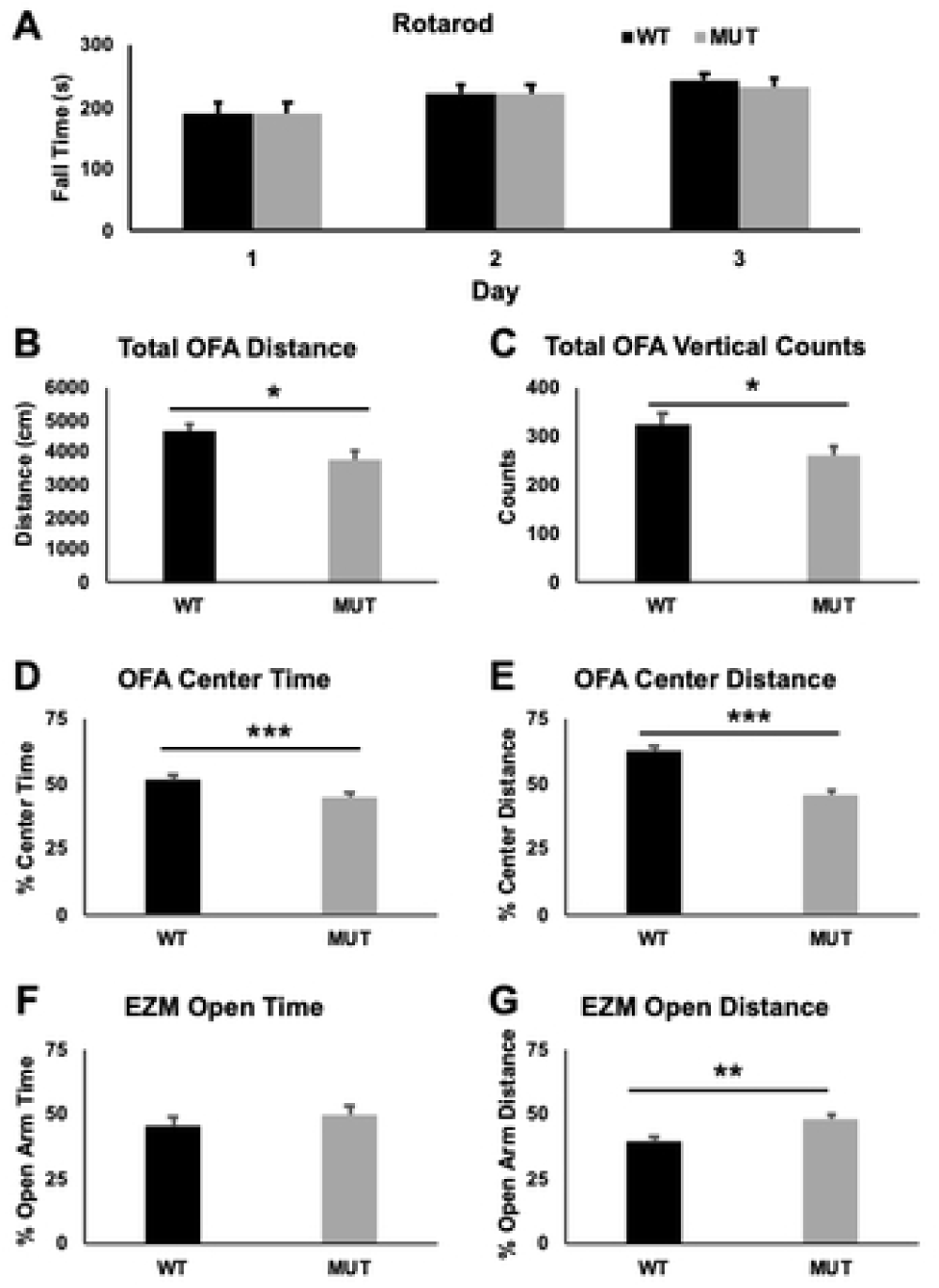
Analysis of motor function and anxiety in *FoxG1*^*MUT/WT*^ mice. **A**) *Foxg1*^*MUT/WT*^ mice (MUT) do not show any changes on the accelerating rotating rod task compared to wild-type (WT) littermate controls (15 weeks old, n=15 for each genotype). **B-E:** Open Field Assay (OFA, 8-11 weeks, WT n=26, MUT n=31). MUT animals have decreased overall distance traveled (**B**, F[1,55]=7.05, p=0.02) and vertical counts (**C**, F[1,55]=4.95, p=0.03). MUT animals spend less percentage time in center area (**D**, F[1,55]=7.233, p=0.009) and travel less percentage difference (**E**, F[1,55]=54.925, p<0.0001) in OFA. **F-G:** Elevated Zero Maze (EZM, 17-22wks, WT n=11, MUT=14). MUT animals did not show any difference in the percent time spent in the open arm (**F**, F[1,23]=1.09, p=0.307) but traveled more percentage distance in the open arm (**G**, F[1,23]=10.576, p=0.004). *p<0.05, *p<0.01, ***P<0.001. No genotype*sex interaction was observed for any measure so only genotype effects were analyzed.

### Heterozygous Foxg1 mice have changes in gait

To further evaluate locomotor function in the MUT animals we conducted gait analysis using a treadmill system. MUT animals showed changes in a number of gait parameters (Table 1), notably decreased forepaw and hindpaw print area, increased percentage of time in forepaw and hindpaw swing, increased hindpaw stride length, decreased rear track width, decreased hindpaw homologous and homolateral coupling, and increased body rotation.

**Table 1:** *Foxg1*^*MUT/WT*^ mice have a variety of changes in gait parameters. Animals were evaluated at 9-12 weeks of life. Two-way ANOVA did not identify any significant genotype*sex interaction effect, so only genotype effects were analyzed. WT n=30, *FoxG1*^*MUT/WT*^ (MUT) n=26. Significant differences are identified with *p<0.05.

| Variable | Genotype | Mean | SEM | F (1,54) | Sig. |
| --- | --- | --- | --- | --- | --- |
| Forepaw Stride (ms) | WT | 280.90 | 5.77 | 2.762 | 0.103 |
|  | MUT | 267.76 | 5.40 |  |  |
| Forepaw % Swing | WT | 0.49 | 0.01 | 7.036 | 0.011* |
|  | MUT | 0.52 | 0.01 |  |  |
| Forepaw Stride Length (mm) | WT | 58.83 | 0.71 | 1.922 | 0.172 |
|  | MUT | 60.18 | 0.66 |  |  |
| Forepaw Print Area | WT | 153.03 | 5.54 | 6.512 | 0.014* |
|  | MUT | 133.67 | 5.19 |  |  |
| Hindpaw Stride (ms) | WT | 284.42 | 5.86 | 0.025 | 0.876 |
|  | MUT | 283.16 | 5.49 |  |  |
| Hindpaw % Swing | WT | 0.53 | 0.01 | 11.629 | 0.001* |
|  | MUT | 0.57 | 0.01 |  |  |
| Hindpaw Stride Length (mm) | WT | 60.37 | 1.03 | 5.574 | 0.022* |
|  | MUT | 63.70 | 0.96 |  |  |
| Hindpaw Print Area | WT | 243.61 | 10.22 | 4.194 | 0.046* |
|  | MUT | 214.92 | 9.58 |  |  |
| Front Track Width (mm) | WT | 11.92 | 0.19 | 0.044 | 0.835 |
|  | MUT | 11.87 | 0.18 |  |  |
| Rear Track Width (mm) | WT | 20.96 | 0.26 | 4.323 | 0.043* |
|  | MUT | 20.24 | 0.24 |  |  |
| Overall Avg Run Speed (mm/s): | WT | 178.64 | 3.76 | 1.184 | 0.282 |
|  | MUT | 184.24 | 3.52 |  |  |
| Forepaw Homolateral Coupling | WT | 0.53 | 0.01 | 3.518 | 0.066 |
|  | MUT | 0.54 | 0.01 |  |  |
| Forepaw Homologous Coupling | WT | 0.50 | 0.00 | 0.468 | 0.497 |
|  | MUT | 0.49 | 0.00 |  |  |
| Forepaw Diagonal Coupling | WT | 0.06 | 0.01 | 1.561 | 0.217 |
|  | MUT | 0.07 | 0.01 |  |  |
| Hindpaw Homolateral Coupling | WT | 0.46 | 0.01 | 6.803 | 0.012* |
|  | MUT | 0.43 | 0.01 |  |  |
| Hindpaw Homologous Coupling | WT | 0.49 | 0.00 | 5.380 | 0.024* |
|  | MUT | 0.48 | 0.00 |  |  |
| Hindpaw Diagonal Coupling | WT | -0.02 | 0.01 | 0.515 | 0.476 |
|  | MUT | -0.03 | 0.01 |  |  |
| Forepaw Gait Angle | WT | 51.99 | 1.13 | 0.178 | 0.675 |
|  | MUT | 52.64 | 1.06 |  |  |
| Hindpaw Gait Angle | WT | 64.63 | 1.28 | 0.370 | 0.546 |
|  | MUT | 63.56 | 1.20 |  |  |
| Body Rotation (deg) | WT | -0.55 | 0.50 | 10.580 | 0.002* |
|  | MUT | -2.76 | 0.46 |  |  |
| Longitudinal Position (mm) | WT | 87.74 | 3.29 | 0.447 | 0.507 |
|  | MUT | 90.75 | 3.08 |  |  |
| Lateral Position (mm) | WT | 82.69 | 0.43 | 0.122 | 0.728 |
|  | MUT | 82.49 | 0.40 |  |  |

### Heterozygous Foxg1 mice have learning and memory deficits

Individuals with FS are considered to have intellectual disability, and previous work found changes in learning and memory in mice expressing a different *Foxg1* allele (16). To evaluate learning and memory in our MUT mice, we performed the conditioned fear task (Figure 3). MUT mice showed increased freezing at baseline (before the conditioned and unconditioned stimulus are presented on day 1). To account for this increased baseline freezing in the MUT animals, we subtracted the amount of freezing in the subsequent context and cue test for analysis. MUT mice showed decreased freezing (over baseline) in either the context (Figure 3B) or cue (Figure 3C). Notably, even when not accounting for the increased baseline freezing observed in MUT animals they still showed decreased freezing both in the context (F[1,28]=5.19, p=0.031) or the cue (F[1,28]=7.66, p=0.01).

**Figure 3.**
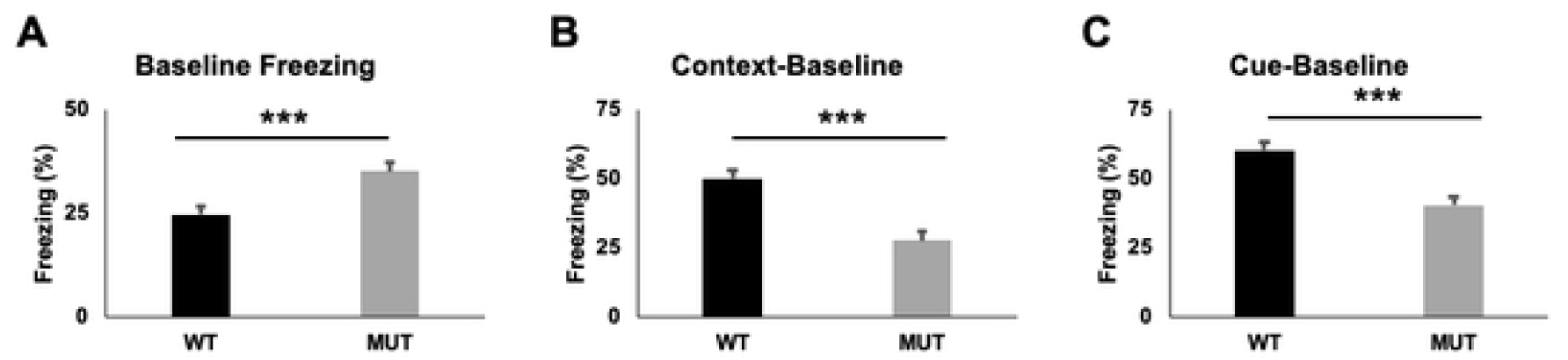
*Foxg1*^*MUT/WT*^ mice show impaired learning on the conditioned fear task. **A**) *FoxG1*^*MUT/WT*^ mice (MUT, n=15) show increased baseline freezing (on training day before stimulus) compared to WT mice (n=15) at 13-14 weeks old (F[1.28]=12.80, p=0.001). **B**) MUT mice have decreased freezing (over baseline freezing) to context (F[1.28]=24.42, p<0.001). **C**) MUT mice have decreased freezing (over baseline freezing) to cue (F[1.28]=25.39, p<0.001). MUT animals also have decreased percentage of absolute freezing in both context (p=0.031) and cue (p=0.01) when not accounting for increased baseline freezing (not shown). No sex or sex*genotype effect observed. ***p<0.001

### Heterozygous Foxg1 mice have alterations in social behavior

To evaluate social and other neuropsychiatric phenotypes in these MUT mice, we performed a battery of tests to assess obsessive compulsive, aggressive, and social behavior. MUT animals buried less marbles that litter-mate WT controls (Figure 4A), indicating decreased obsessive-compulsive features. It was noted that MUT animals seemed more aggressive, with increased biting of human handlers and instances of fighting with home-cage littermates. We evaluated social dominance with the Tube Test, exposing each animal to two challenges over two days. MUT animals had a higher average rate of winning at the tube test than WT animals (Figure 4B). In the Nest Building task, MUT animals build lower quality nests (Figure 4C) and had a smaller amount of the nestlet shredded for each nest (Figure 4D).

**Figure 4.**
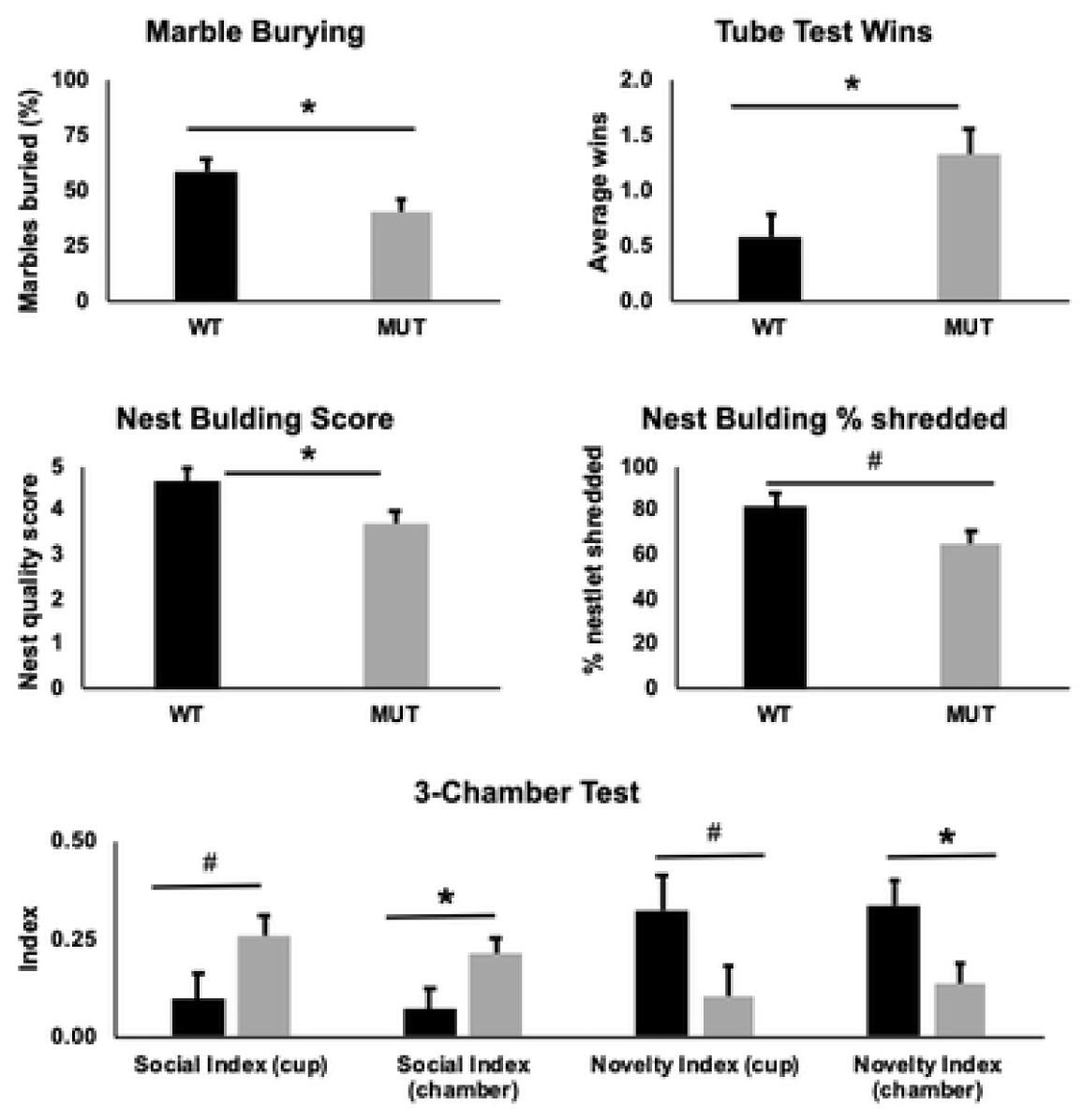
*Foxg1*^*MUT/WT*^ mice show changes in compulsive and social behavior tests. **A**) *Foxg1*^*MUT/WT*^ mice (MUT, n=15) have a decreased percentage of marbles buried in the marble burying task compared to WT (n=15) at 13 weeks (F[1.28]=4.89, p=0.035). **B**) MUT (n=15) mice show increased average number of wins (over 2 bouts) compared to WT (n=15) at 11-12 weeks (F[1,28]=5.965, p=0.021). At 14-15 weeks old, MUT (n=15) animals had a decreased next building score (**C**, F[1,28]=7.18, p=0.012) and trended towards a decreased percentage of nestlet shredding (**D**, F[1,28]=4.165, p=0.051) compared to WT (n=15) animals. **E**) MUT mice (n=16) have changes in social behavior on the 3-chamber task at 9 weeks compared to WT (n=11) mice. MUT animals trended to have increased Social Index in close proximity to cup with stranger mouse (F[1,25]=3.54, p=0.072) and significant increase in Social Index of time in chamber with stranger mouse (F[1,25]=4.66, p=0.041). In contrast, MUT mice spent more time with a familiar mouse than a new stranger mouse (Novelty Index), with a trend towards decreased Novelty Index to cup (F[1,25]=3.55, p=0.054) and significant decreased Novelty Index in the chamber (F[1,25]=6.05, p=0.021). No sex*genotype effect observed for any measures. #p<0.1, *p<0.05

To evaluate social interaction, we evaluated the MUT mice using the 3-Chamber task. On the first day, the amount of time interacting with a cup containing a mouse compared to an empty cup is evaluated. MUT mice showed a trend towards increased Social Index using time interacting closely with the cup containing the mouse (Figure 4E, Social Index Cup) and significant increased Social Index using time within the chamber with the cup containing the mouse (Figure 4E, Social Index Chamber). On the subsequent day, social novelty was assessed by exposing the test mouse to a cup containing the mouse from the previous day (Familiar) versus a cup containing a new mouse (Novel) to assess the interest in social novelty. MUT mice showed a trend towards decreased interest in the novel mouse cup (Figure 4E, Novelty Index Cup) and significant difference in the interest in the chamber containing the novel mouse (Figure 4E, Novelty Index Chamber).

### Imaging revealed decreased overall and regional brain volumes in heterozygous Foxg1 mice

#### Whole Brain Volume

As expected, the results of the t-test with WBV revealed that the WBVs of the HET mice were smaller than those of the WT group (*p*=.00662). See Table 2.

**Table 2:** Volumetric assessment of *Foxg1*^*MUT/WT*^ mice using MRI. Animals were assessed at ∼48 wks of life, n=6 for each genotype. p-values that were below threshold for Bonferroni multiple testing correction are shown in bold.

| ROI | Group | Mean-no WBV | SD-no WBV | p-value no WBV | Mean-WBV Corrected | SD-WBV Corrected | p-value<sup>#</sup> |
| --- | --- | --- | --- | --- | --- | --- | --- |
| Whole Brain Volume (WBV) | WT | 150.8333 | 4.92875 | **\*.00662** | --- | --- | --- |
|  | MUT | 138.1869 | 7.2541 |  |  |  |  |
| Lateral Ventricle | WT | 1.65265 | 0.20841 | **\*.00578** | 0.01095 | 0.0013 | 0.02207 |
|  | MUT | 1.27545 | 0.05663 |  | 0.00925 | 0.00057 |  |
| Intraventricular Foramen | WT | 0.0039 | 0.00306 | 0.14837 | 0.00003 | 0.00002 | 0.1668 |
|  | MUT | 0.00173 | 0.00093 |  | 0.00001 | 0.00001 |  |
| Hippocampus | WT | 6.47802 | 0.18381 | **\*.00371** | 0.04297 | 0.00126 | 0.70867 |
|  | MUT | 5.90553 | 0.29944 |  | 0.04274 | 0.00066 |  |
| Olfactory System | WT | 24.58858 | 1.01571 | **\*.00226** | 0.16316 | 0.0084 | 0.63568 |
|  | MUT | 22.28573 | 0.94077 |  | 0.16136 | 0.00275 |  |
| Basal Ganglia | WT | 9.13177 | 0.58828 | **\*.00425** | 0.0605 | 0.0024 | 0.0478 |
|  | MUT | 7.98298 | 0.47336 |  | 0.05777 | 0.00167 |  |
| Olfactory Nerve | WT | 0.00073 | 0.00054 | 0.55723 | 0 | 0 | 0.76566 |
|  | MUT | 0.00057 | 0.0004 |  | 0 | 0 |  |
| Corpus Callosum | WT | 12.3984 | 0.42803 | **\*.00174** | 0.08221 | 0.00148 | **\*.00109** |
|  | MUT | 10.65882 | 0.79888 |  | 0.07707 | 0.00219 |  |
| Anterior Commissure Olfactory Limb | WT | 0.64525 | 0.04162 | 0.09611 | 0.00428 | 0.00029 | 0.79429 |
|  | MUT | 0.58655 | 0.06466 |  | 0.00424 | 0.00029 |  |
| Lateral Olfactory Tract | WT | 1.36563 | 0.08943 | 0.04843 | 0.00907 | 0.00079 | 0.45808 |
|  | MUT | 1.20957 | 0.13989 |  | 0.00874 | 0.0007 |  |
| Internal Capsule | WT | 3.91435 | 0.19739 | **\*.00437** | 0.02594 | 0.00058 | 0.01408 |
|  | MUT | 3.46397 | 0.22565 |  | 0.02505 | 0.00043 |  |
| Anterior Commissure Temp Limb | WT | 0.47452 | 0.00716 | 0.27234 | 0.00315 | 0.00007 | **\*.00121** |
|  | MUT | 0.46288 | 0.02247 |  | 0.00335 | 0.00008 |  |
| Stria Terminalis | WT | 0.00108 | 0.00091 | 0.47036 | 0.00001 | 0.00001 | 0.35297 |
|  | MUT | 0.00142 | 0.00058 |  | 0.00001 | **0** |  |
| Amygdala | WT | 0.8005 | 0.02179 | **\*.00066** | 0.00531 | 0.00012 | **\*.00292** |
|  | MUT | 0.6532 | 0.05519 |  | 0.00473 | 0.00029 |  |
| Hypothalamus | WT | 9.64162 | 0.3497 | 0.14864 | 0.06392 | 0.001 | 0.00811 |
|  | MUT | 9.25945 | 0.47951 |  | 0.06703 | 0.00189 |  |
| Thalamus | WT | 18.01777 | 0.67212 | **\*.00630** | 0.11945 | 0.00207 | 0.27956 |
|  | MUT | 16.30582 | 0.97022 |  | 0.11798 | 0.00238 |  |
| Fasciculus Retroflexus | WT | 0.07788 | 0.00882 | 0.09664 | 0.00052 | 0.00005 | 0.79877 |
|  | MUT | 0.07015 | 0.00471 |  | 0.00051 | 0.00004 |  |
| Stria Medullaris | WT | 0.29677 | 0.01387 | 0.02382 | 0.00197 | 0.00006 | 0.39622 |
|  | MUT | 0.26662 | 0.02284 |  | 0.00193 | 0.00009 |  |
| Midbrain | WT | 14.497 | 0.67974 | **\*.00619** | 0.0961 | 0.00265 | 0.84772 |
|  | MUT | 13.31745 | 0.38658 |  | 0.09656 | 0.00508 |  |
| Hindbrain | WT | 0.00082 | 0.00056 | 0.67908 | 0.00001 | 0 | 0.79939 |
|  | MUT | 0.00068 | 0.00052 |  | 0 | 0 |  |
| Pons | WT | 26.39558 | 1.07066 | 0.07458 | 0.17496 | 0.00172 | **\*.00049** |
|  | MUT | 25.12625 | 1.1367 |  | 0.1819 | 0.00262 |  |
| Medulla | WT | 14.43622 | 1.172 | 0.32998 | 0.09564 | 0.0059 | 0.49559 |
|  | MUT | 13.6141 | 1.56915 |  | 0.09827 | 0.00693 |  |
| Fornix System | WT | 4.49635 | 0.24178 | 0.07291 | 0.02981 | 0.00136 | 0.41149 |
|  | MUT | 4.20985 | 0.25323 |  | 0.03048 | 0.00133 |  |
| Pituitary | WT | 1.51792 | 0.2617 | 0.94945 | 0.01005 | 0.00158 | 0.3395 |
|  | MUT | 1.52813 | 0.28226 |  | 0.01102 | 0.00176 |  |

<sup>#</sup>controlling for WBV; \*FDR corrected, $p < .05$

#### Regions of Interest

FDR-corrected findings of the t-tests revealed that out of the 23 ROIs, nine reached statistical significance: Lateral Ventricle, Hippocampus, Olfactory System, Basal Ganglia, Corpus Callosum, Internal Capsule, Amygdala, Thalamus, and Midbrain. All the aforementioned structures showed significantly smaller volumes in the HET as compared to WT mice (Table 2).

#### Regions of Interest (corrected for WBV)

FDR-corrected findings of the t-tests revealed that out of the 23 ROIs corrected for WBV (i.e., the relative differences in brain structures after total brain volume has been equated across groups), four ROIs reached statistical significance: Corpus Callosum, Amygdala, Anterior Commissure Temp Limb, and Pons. The Corpus Callosum and Amygdala remained smaller for HET than WT mice, suggesting a marked reduction in volume for these areas even after accounting for the globally reduced brain volume associated with the HET mice. In contrast, interestingly, the Anterior Commissure Temp Limb and the Pons were relatively *larger* for HET than WT mice, suggesting that relative to their overall smaller brain volumes, HET mice show larger than expected volumes in these two areas (Table 2).

## Discussion

Recent work in neurodevelopmental disorders such as Rett syndrome has identified potential for symptomatic reversal when the missing protein function is restored, even after symptom onset, leading to the development of gene therapy or other methods to correct the genetic deficit. While FS is associated with structural brain changes originating during embryogenesis, the ongoing expression of FoxG1 in the adult brain and development of phenotypes when FoxG1 function is removed postnatally provides hope that a postnatal window of opportunity exists for meaningful interventions that could modify the overall disease course. Here we created and characterized a new model of FS and identified phenotypes that model aspects of the human disease and others that have not been reported in humans with FS. As has been found in other mice with heterozygous mutations in *Foxg1* (summarized in Table 3), we observed structural brain abnormalities including an overall decreased brain size as well as decreased size of the corpus callosum, even after correcting for the decreased whole brain volume. Unexpectedly, when corrected for whole brain volume, the pons was relatively larger in the heterozygous mutant animals compared to littermate controls. This likely reflects the role of FoxG1 in telencephalon development, resulting in an overall decrease in cortex and forebrain structures with preservation of hindbrain structures. As can be seen in Table 3, the findings presented in this manuscript are broadly similar to those previously reported in other heterozygous mouse models of FS.

**Table 3:**
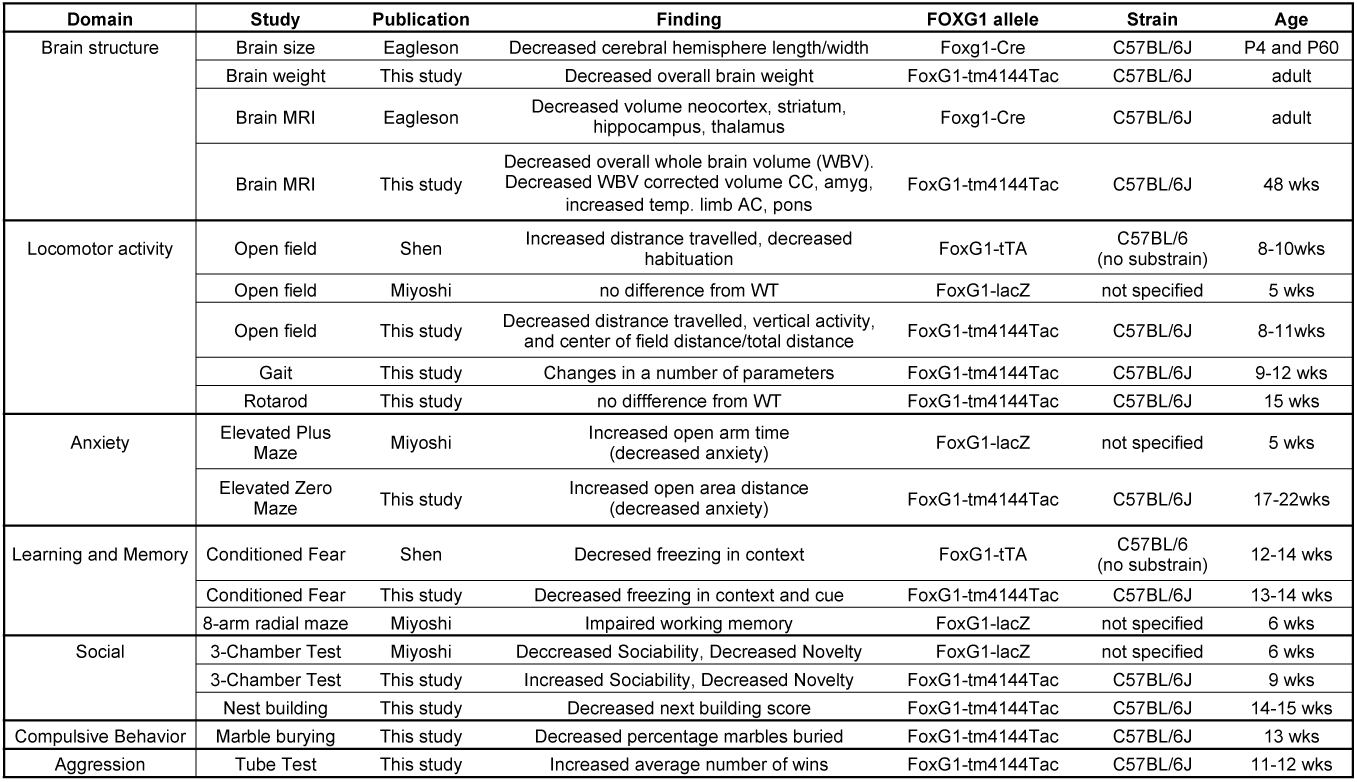
Summary and comparison of phenotypes observed to previous published findings. CC=corpus callosum, AC=anterior commissure, amyg=amygdala. References: Eagleson (15), Shen (16), Miyoshi (14)

Gross motor skills are markedly impaired in people with FS, and here we found that heterozygous mutant mice showed overall hypoactivity compared to wild-type littermate controls. Previous published work has found either no change in locomotor activity, or overall hyperactivity and a failure to decrease activity overtime in mutant animals (See Table 3). While these results seem to be incongruous, aspects of the different experiments make it difficult to directly compare. Each experiment used a different allele of *Foxg1*, and while each allele should completely disrupt FoxG1 protein production, subtle differences in the alleles could manifest as different behaviors. More importantly, different ages were characterized in each experiment, which could markedly influence overall locomotor activity. Longitudinal characterization of this phenotype will be needed to determine if there is an evolving pattern of locomotor behavior. Finally, the lack of clarity regarding the background strains used in previously published work also makes direct comparisons difficult, as background strain can have marked effects on genetically based phenotypes. In addition to the characterization of locomotor activity, here we also characterized gait and found a number of changes. This represents a clinically meaningful phenotypes, as gait is markedly disrupted or absent in people with FS (3). Unexpected, the heterozygous mutant animals did not show any differences in motor coordination or motor learning on the rotarod task. Future work exploring other motor functions is needed to fully characterize the spectrum of phenotypes present in this important domain.

In this work we found clear impairment in learning and memory as assessed by the conditioned fear task. This is partially consistent with changes observed in a different *Foxg1* allele by Shen *et. al* (16), however they only observed changes in the context and not the cue stimulus. While both studies were conducted at roughly the same age, the caveats regarding specific allele and strain mentioned above remain. A recent study identified deficits in working memory using the 8-arm radial maze in a different allele of *Foxg1* (14). Together these results across different *Foxg1* alleles at different ages indicate that deficits in learning and memory are a consistent feature of heterozygous mutations in *Foxg1*, representative of the intellectual disability seen in people with FS (3). This deficit in learning and memory may be one of the clinical features with the greatest potential for modification with post-natal restoration of FoxG1 function, as previous work has demonstrated that removal of FoxG1 function specifically from excitatory neurons at P60 caused deficits in learning and memory as assessed on the Conditioned Fear task and the Morris Water Maze as well as reduced hippocampal long term potentiation (7), and excitingly Miyoshi et al recently demonstrated that transplantation of GABAergic precursor cells at P7 can rescue the working memory deficit in *Foxg1* heterozygous animals (14).

Consistent with published work(14), we found decreased interest to a novel mouse in the 3-Chamber test, however in contrast we found increased rather than decreased sociability in the same task (Table 3). These experiments were conducted at different ages and future longitudinal studies will be needed to determine if this social interaction difference is age related or a function of different *Foxg1* alleles or background strains. As autistic features such as poor social interaction and eye contact are present in FS (3), these abnormal social behaviors are again important clinically relevant phenotypes that also can be modified with GABAergic precursor cell transplantation (14). We also found additional behavioral changes, such as decreased anxiety, poor nest building, decreased compulsive stereotypical behavior, and increased aggression. While these are not clearly reported clinical issues in people with FS, they represent additional phenotypes that can be evaluated in preclinical therapy evaluation.

This newly generated allele was developed by the incorporation of a “STOP” cassette containing a splice acceptor and multiple translational stop codons flanked by loxP sequences inserted upstream of the coding sequence of *Foxg1*, with the goal of disrupting expression of FoxG1 protein while retaining the ability to restore expression by the Cre-dependent removal of the STOP cassette. A similar strategy was successfully utilized to demonstrate the ability for post-symptomatic rescue of Rett syndrome mice (8). Future work will focus on the evaluation of the ability of this newly generated allele to be “rescued” by exposure to the Cre recombinase. Although this strategy has been successful in Rett syndrome, differences exist that may prevent this new allele to work as effectively as in that case. Whereas in Rett syndrome the STOP cassette was introduced into an exon between two coding introns, in mice the coding sequence of *Foxg1* is contained entirely within a single exon that is preceded by a non-coding exon. Although we have demonstrated here that this STOP cassette does disrupt the production of FoxG1 protein, there exists the potential that even after Cre recombination FoxG1 protein remains disrupted. An alternative approach to postnatal restoration of FoxG1 protein has been developed recently by Miyoshi et al (14) using a tetracycline-dependent transgenic system to drive expression of a *Foxg1* transgene. In combination with a *Foxg1* heterozygous mutant allele, postnatal rescue of phenotypes could be evaluated with such a system. Regardless of the exact method, it is crucial to evaluate the potential to modify clinical features in FS by post-natal restoration of FoxG1 function to determine if strategies such as gene therapy are worthwhile to pursue, or if alternative strategies should be developed to treat this severe neurodevelopmental disorder.

## Acknowledgments

We thank Chris Svitek for technical assistance. The content is solely the responsibility of the authors and does not necessarily represent the official views of the National Institutes of Health or the Eunice Kennedy Shriver Child Health and Human Development Institute (NICHD).

